# Adapting in larger numbers can increase the vulnerability of *Escherichia coli* populations to environmental changes

**DOI:** 10.1101/546119

**Authors:** Yashraj Chavhan, Shraddha Karve, Sutirth Dey

**Affiliations:** Indian Institute of Science Education and Research (IISER) Pune, Dr. Homi Bhabha Road, Pashan, Pune, Maharashtra, 411008, India.; Department of Evolutionary Biology and Environmental Studies, University of Zurich, Winterthurerstrasse 190, 8057 Zurich, Switzerland

**Keywords:** population size, adaptation speed, efflux activity, character decay

## Abstract

Larger populations generally adapt faster to their existing environment. However, it is unknown if the population size experienced during evolution influences the ability to face sudden environmental changes. To investigate this issue, we subjected replicate *Escherichia coli* populations of different sizes to experimental evolution in an environment containing a cocktail of three antibiotics. In this environment, the ability to actively efflux molecules outside the cell is expected to be a major fitness-affecting trait. We found that all the populations eventually reached similar fitness in the antibiotic cocktail despite adapting at different speeds, with the larger populations adapting faster. Surprisingly, whereas efflux activity enhanced in the smaller populations, it decayed in the larger ones. The evolution of efflux activity was largely shaped by pleiotropic responses to selection and not by drift. This demonstrates that quantitative differences in population size can lead to qualitative differences (decay/enhancement) in the fate of a character during adaptation to identical environments. Furthermore, the larger populations showed inferior fitness upon sudden exposure to several alternative stressful environments. These observations provide a novel link between population size and vulnerability to environmental changes. Counter-intuitively, adapting in larger numbers can render bacterial populations more vulnerable to abrupt environmental changes.

## Introduction

Population size is a key ecological parameter that influences the rate at which asexual populations evolve (Gerrish and Lenski 1998; Wilke 2004; Desai and Fisher 2007; Desai et al. 2007). All else being equal, larger populations are supposed to evolve faster as they are expected to have access to greater variation (Orr 2000; Wilke 2004; Desai and Fisher 2007; Desai et al. 2007; Sniegowski and Gerrish 2010). Moreover, the efficiency of natural selection, in favoring beneficial mutations and keeping out deleterious ones, increases with increasing population size (Petit and Barbadilla 2009; Chavhan et al. 2019), which is also expected to increase the rate of adaptation. However, little is known about how evolving large asexual populations fare when their environment changes abruptly. Are their performances comparable with smaller populations that have evolved in the same environment?

Consider a clonally derived large asexual population that has evolved in a constant environment for an extended period. The ability of such a population to face sudden environmental changes would be determined by the variation accumulated during evolution in the constant environment. However, the population size experienced during evolution will influence variation in two contrasting ways. On the one hand, larger asexual populations are expected to stumble upon more mutations during adaptation (Desai and Fisher 2007; Desai et al. 2007; Sniegowski and Gerrish 2010). On the other hand, since natural selection is more efficient in larger populations, it can lead to a rapid increase in the average fitness and severe reduction in the genetic variation of such populations (Desai and Fisher 2007; Sniegowski and Gerrish 2010; Chavhan et al. 2019). Such reduction in variation can potentially be detrimental if the environment changes suddenly, particularly if high fitness in the old environment is correlated with low fitness in the new one (antagonistic pleiotropy *sensu* Cooper and Lenski (2000); Cooper 2014). Thus, the actual amount of variation available to the population would be determined by an interaction between these two opposing aspects.

Asexual populations of very different sizes also have markedly different accessibilities to beneficial mutations in identical environments (Wilke 2004; Desai and Fisher 2007; Sniegowski and Gerrish 2010; Chavhan et al. 2019). This is because beneficial mutations that confer higher fitness gains are generally rarer (Kassen and Bataillon 2006; Eyre-Walker and Keightley 2007; Perfeito et al. 2007; Sniegowski and Gerrish 2010; Neher 2013)). Consequently, whereas adaptation in very large populations is driven predominantly by rare large-effect beneficial mutations, small populations typically adapt via relatively common small-effect beneficial mutations (Sniegowski and Gerrish 2010). The ability of a small population to face environmental changes is also expected to be different from that of a larger one. This notion stems primarily from theoretical studies which predict that the pleiotropic effects of large- and small-effect beneficial mutations should be very different (Lande 1983; Orr and Coyne 1992). For example, Lande’s model for studying the response to selection on beneficial mutations of varying sizes assumed that major beneficial mutations have substantial pleiotropic costs while minor beneficial mutations have none (Lande 1983). It has been suggested that a better assumption would be that the deleterious pleiotropic effects of a beneficial mutation are proportional to the size of the benefit it confers (Orr and Coyne 1992). Furthermore, recent empirical investigations have found that deleterious mutations that confer larger fitness deficits also tend to have more pleiotropic effects (Cooper et al. 2007). Overall, the extant literature suggests larger beneficial mutations may have greater deleterious pleiotropic effects. Given that large populations adapt primarily via rare large-effect mutations and small populations via relatively common mutations of small effect (Sniegowski and Gerrish 2010), it is expected that larger populations would suffer heavier pleiotropic disadvantages. Thus, if asexual populations of different sizes adapt to the same constant environment for an extended period, larger populations can become inferior to smaller ones in terms of their immediate response to environmental changes.

In this study, we used experimental evolution to examine the above notion. Specifically, we propagated replicate *Escherichia coli* populations of different sizes in a constant environment for ∼ 380 generations. This constant environment contained an unchanging sub-lethal cocktail of three antibiotics, namely, norfloxacin, rifampicin, and streptomycin. When all the populations reached similar fitness in this environment, we estimated their ability to face sudden changes in environmental conditions using two different approaches. First, we studied the evolution of energy-dependent efflux activity (EA), which represents the generic capacity of bacteria to actively transport unwanted molecules out of their cells, and is a critical component of xenobiotic metabolism (Sun et al. 2014). EA is known to be one of the broad-based mechanisms in bacteria for fighting multiple stresses including antibiotics (Kumar and Schweizer 2005), heavy metals (Nies 2003; Poole 2005), bile salts (Thanassi et al. 1997), organic solvents (Fernandes et al. 2003), intercalating mutagens (Ma et al. 1993; Nishino et al. 2009), etc. This makes EA a good candidate character to study the ability of bacterial populations to thrive in the face of sudden environmental stress (Karve et al. 2015). Second, we directly tested the fitness of our populations in several alternative environments which are known to affect *E. coli* very differently as compared to the three antibiotics in the selection environment.

The three antibiotics used in our selection environment had very different mechanisms and sites of action (Drlica and Zhao 1997; Campbell et al. 2001; Sharma et al. 2007). Evolution in this environment is expected to favour the presence of EA. However, we found that whereas larger populations undergoing fast adaptation experienced decay of EA, smaller populations undergoing slow adaptation experienced enhanced EA. These results were attributable to correlated responses to selection rather than the accumulation of contextually neutral mutations via genetic drift. The larger population also had lower fitness upon exposure to four different alternative environments. This demonstrates that highly efficient selection during rapid adaptation in large populations can render them vulnerable in terms of their response to environmental changes.

Adaptation to a given environment is expected to result either in enhancement/maintenance or in decay of a biological character (but not both). To the best of our knowledge, this is the first study to show that even in the absence of major effects of drift, a biological character under selection can decay or enhance depending on the size of the adapting populations.

## Materials and methods

### Experimental evolution and measurement of adaptive dynamics

The maintenance protocol of these selection lines have been previously described in another study (Chavhan et al. 2019). We derived 24 microbial populations from a single *Escherichia coli* MG 1655 colony and randomly distributed them among three population size treatments, namely LL, SL, and SS (refer to the next paragraph for the details of this nomenclature), leading to 8 independently evolving replicate populations per treatment. The populations evolved in a constant environment made of nutrient broth containing a sub-lethal cocktail of three antibiotics (henceforth called ‘selection environment’) under batch culture for ∼380 generations (see Supplementary Methods for the detailed composition of the nutrient broth). The three antibiotics used were norfloxacin (0.015 μg/ml), rifampicin (6 μg/ml) and streptomycin (0.1 μg/ml). These antibiotics target different cellular mechanisms: norfloxacin interferes with DNA replication (Drlica and Zhao 1997), rifampicin affects RNA transcription (Campbell et al. 2001), while streptomycin affects protein translation (Sharma et al. 2007).

We propagated the three population types at different population sizes. The size of a typical periodically bottlenecked asexual population depends on three interdependent parameters: N_0_ (the number of individuals in the bottleneck), N_f_ (the number of individuals before the bottleneck), and g (the number of generations between successive bottlenecks). Since these populations grow via binary fissions, N_f_ = N_0_ × 2^g^ (Lenski et al. 1991). The conventional measure of size in bottlenecked populations is the harmonic mean of population size (HM = N_0_ × g) (but also see Chavhan et al. (2019) for a measure of population size relevant for predicting the extent of adaptation in such systems). Thus, bottleneck properties are instrumental in shaping the size of such populations. Our experiment had three different population size treatments, called LL, SL, and SS. The first letter of a population type’s name represents a relative measure of the harmonic mean size (L ≈ 3.3 x 10^10^; S ≈ 2.0 x 10^6^) and the second letter represents a relative measure the culture volume (L refers to 100 ml and S refers to 1.5 ml). The density of individuals (number of individuals per unit volume) was identical across the three population types at the beginning of the experiment. Moreover, whereas LL faced lenient bottlenecks (1/10; g = 3.3), the bottleneck ratios were much harsher in SS (1/10^4^; g = 13.3) and SL (1/10^6^); g = 19.9). This ensured that LL >> SL = SS in terms of HM.

We computed the speed of adaptation (SoA) of the three population types using the fitness trajectories over ∼380 generations reported in an earlier study (Chavhan et al. 2019). The fitness trajectories of all three population types had displayed diminishing returns, which is a common observation in evolution experiments with microbes (Cooper and Lenski 2000; Schoustra et al. 2009; Couce and Tenaillon 2015). Therefore, we quantified SoA as the maximum slope of fitness trajectories observed during the evolution experiment. SoA was quantified in terms of two measure of fitness, namely, carrying capacity (K, the maximum optical density reached in a growth assay) and maximum growth rate (R, the maximum slope of the growth curve during the assay).

The experimental design of our study is shown schematically in Fig. 1.

**Fig. 1.**
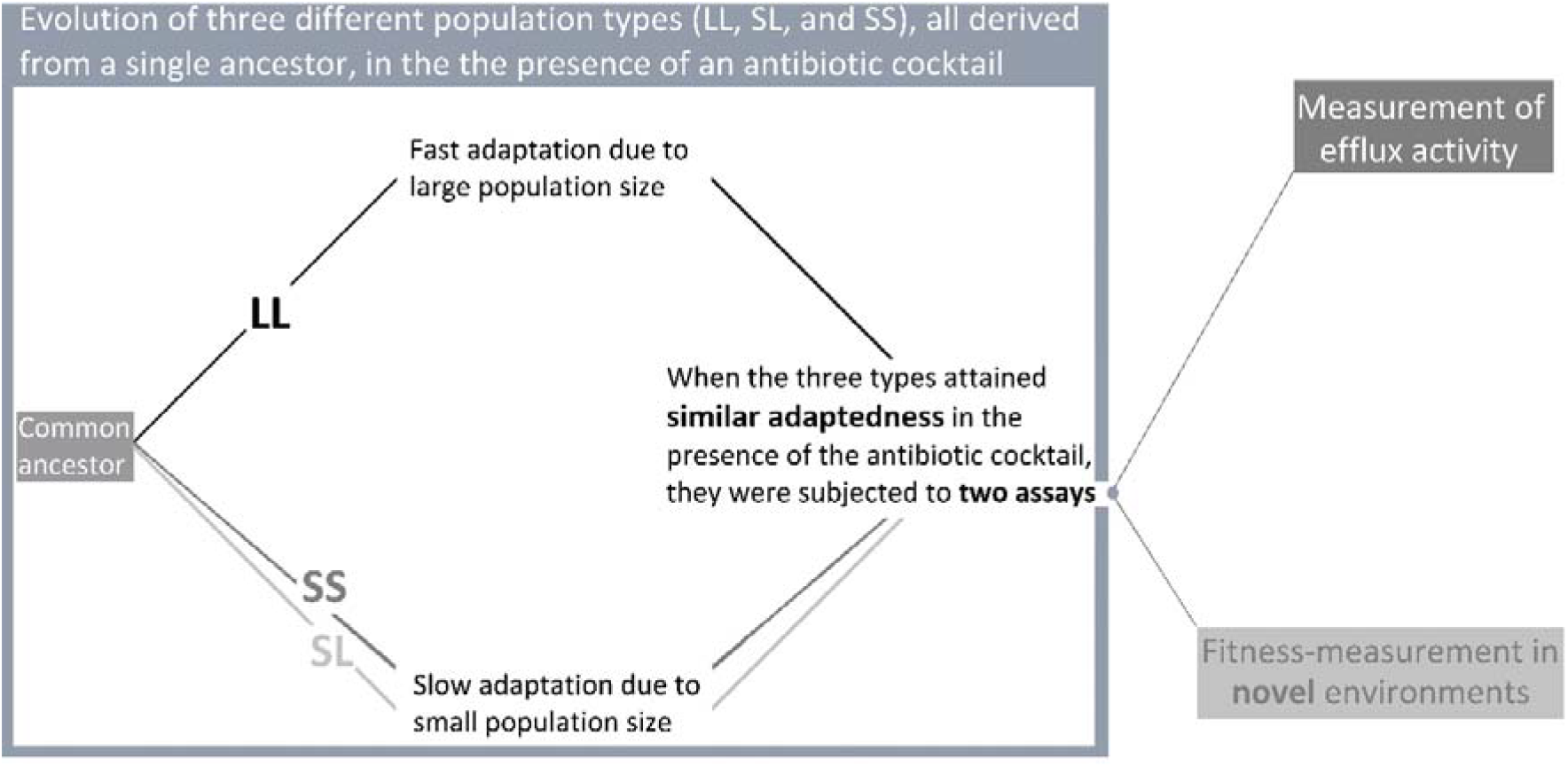
A schematic representation of our study. Note that the assays reported here had been carried out when the three population types (LL, SL, and SS) did not have significantly different fitness in the presence of the antibiotic cocktail, i.e. they were at similar levels of adaptedness.

### Measurement of efflux activity (EA)

We measured the generic EA of the three population types at the beginning and the end of the above evolution experiment using a previously-established protocol (Webber and Coldham 2010; Karve et al. 2015). Specifically, we measured the efficiency with which bacteria could transport a small, foreign molecule out of their cells (see Supplementary Methods).

We used one-sample t-tests to determine if the population types had evolved significantly different EA than the ancestor. We also used a one-way ANOVA with population type (LL, SL, or SS) as the categorical predictor and efflux activity (EA) as the dependent variable to determine the statistical significance of EA differences between the three types. We made sure that the EA assays were carried out when the fitness of the three types were statistically indistinguishable in their selection environment (Fig. 1). Furthermore, all the three population types were derived from a common ancestor. Therefore, the results of these assays can only be attributed to differences in their population sizes. We further used Cohen’s *d* for comparing the significance of differences in the EA of the three population types in terms of effect sizes (Cohen 1988). We also tested if the variation across replicates was significantly different for the three population types. To this end, we compared the variances of LL, SL, and SS lines in a pairwise manner using the Fligner-Killeen test for homogeneity of variances (Fligner and Killeen 1976; Donnelly and Kramer 1999).

### Fitness assays in alternative environments

We quantified the fitness of the three population types in four distinct alternative environments at the end of our evolution experiment (the same time-point at which EA was measured). The design of our study demanded that each alternative environment must impose a challenge that is known to be different from the one imposed by the antibiotic cocktail. Otherwise, the fitness of a population in an alternative environment could be trivially predicted from its fitness in one of the antibiotics in the selection environments. We used Ampicillin as an alternative stress because it has a different site and mechanism than all the three antibiotics used in the cocktail (ampicillin is a β-lactam antibiotic which inhibits cell-wall synthesis) (Waxman and Strominger 1983). Similarly, we used high concentrations of copper (heavy metal stress) as another alternative environment. At high concentrations, the incompletely filled d-orbitals of Cu^2+^ ions form unspecific complex compounds which are toxic to the cellular physiology (Nies 1999). Further, we also used two nutritionally challenging minimal media based on sorbitol and urea as the only carbon sources, respectively.

We revived the end-point cryostocks from our selection-experiment and grew them in nutrient broth (without antibiotics) for 12 hours, which represents ∼ 6.6 doublings. Thus, any lingering physiological effects of stress due to antibiotics were ameliorated. Since all the population types had evolved in the same environment, the effects of the historic environment were not an issue in our study. We carried out automated growth-assays on these populations in the alternative environments using 96-well tissue culture plates in a well-plate reader (Synergy HT, BIOTEK ® Winooski, VT, USA). We used OD at 600 nm as the measure of bacterial density, and assayed growth from each cryostock-derived population in three measurement-replicates. The 96-well-plate was incubated at 37□C and shaken continuously at 150 rpm. The culture volume in each well was 180 μl. The reader took OD readings every 20 minutes, which gave rise to high resolution sigmoidal growth-curves. We used two measures of fitness: (1) Carrying capacity (*K*, the maximum OD achieved during the growth curve) and (2) Growth rate (*R*, the highest slope of growth curve measured over a dynamic window of ten OD readings).

The fitness trends in alternative environments were analyzed in two ways. First, we performed a pooled analysis using a mixed-model ANOVA with population type (three levels: LL, SL, and SS) and alternative environment (four levels: presence of ampicillin, urea as the sole carbon source, sorbitol as the only carbon source, and presence of high [Cu^2+^]) as fixed factors crossed with each other, and replicate number (1-8) as a random factor nested in population type.

Since differences in performance within any one of the four alternative environments could have driven the results of the pooled analysis on their own, we also performed a separate analysis for each alternative environment. Each of these tests involved a nested-design ANOVA with population type (LL, SL, and SS) as the fixed factor, and replicate number (1-8) as a random factor nested in population type. We controlled for false discovery rates (FDR) in these individual ANOVAs using the Benjamini-Hochberg procedure (Benjamini and Hochberg 1995). We used a stringent set of four conditions to determine the significance of the tests done for each environment individually: (1) The ANOVA done over the triplet of LL, SL, and SS should have *P* smaller than the corresponding Benjamini-Hochberg critical value. (2) The difference within the population type triplet has large effect size (partial η^2^ > 0.14) (Cohen 1988). (3) Tukey’s HSD should reveal significant pairwise differences (*P* < 0.05). (4) The pairwise differences should have medium or large effect sizes (Cohen’s *d*). We counted the results of a test as significant only when all four of these conditions were met simultaneously.

## Results

### LL populations had greater speed of adaptation (SoA)

There was a significant effect of population size on SoA, both in terms of K (Fig. 2a. *P* = 1.505×10^-8^; F_(2,21)_ = 47.870) and R (Fig. 2b. *P* = 2.73×10^-6^; F_(2,21)_ = 25.070) (one-way ANOVA, N = 8). Tukey’s HSD (pair-wise post-hoc) suggested the following relationships for K: LL > SL (*P* = 1.501×10^-4^), LL > SS (*P* = 1.403×10^-4^), SL > SS (*P* = 0.004), and R: LL > SL (*P* = 1.55×10^-4^), LL > SS (*P* = 1.45×10^-4^) and SL ∼ SS: (*P* = 0.907). Taken together, this suggests that LL populations adapted faster than both SL and SS.

**Fig. 2.**
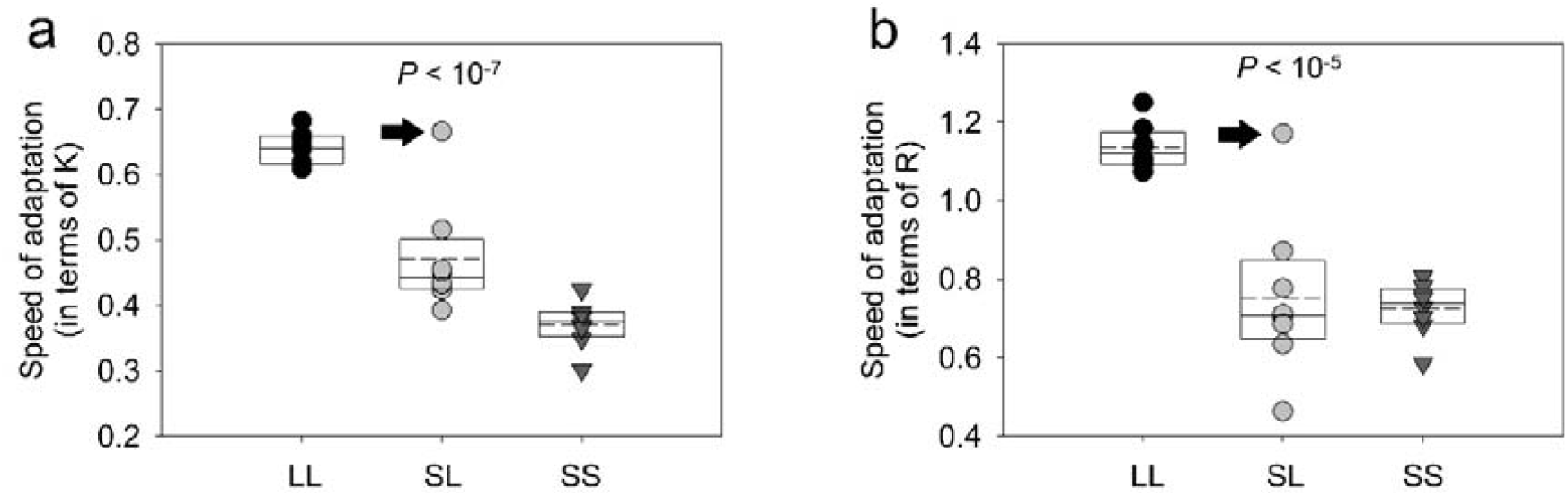
Speed of adaptation during evolution to the antibiotic cocktail. The solid lines in the box-plots mark the 25^th^, 50^th^, and 75^th^ percentiles; the dashed lines within the box-plots represent means (N=8). (a) Speed of adaptation in terms of K. (b) Speed of adaptation in terms of R. The grey data points marked with an arrow represent the only non-LL population that lost the ancestral efflux activity (see the text for details). Note that it was the same replicate population which was an outlier in terms of both K and R in the SL treatment.

### LL populations evolved reduced efflux activity (EA)

We found that whereas LL lost the efflux activity (EA) with respect to the common ancestor, SL and SS gained it (Fig. 3a) (one sample t-test against the ancestral efflux activity: *P =* 0.003086, Cohen’s *d* = 1.26 (large effect) (LL); *P* = 0.029800, Cohen’s *d* = 1.25 (large effect) (SL); *P =* 0.000823, Cohen’s *d* = 2.28 (large effect) (SS). One-way ANOVA (N=8) across the three population types revealed a significant main effect of population type (*P* = 2.055×10^-6^; F_(2,21)_ = 26.044) and Tukey’s HSD for the pairwise comparisons showed LL < SL (*P* = 2.710×10^-4^), LL < SS (*P* = 1.411×10^-4^) and SL ∼ SS (*P* = 0.155). Furthermore, the statistically significant pairwise differences (LL-SL and LL-SS) also had very large effect sizes: Cohen’s *d* = 2.512 for LL-SL; Cohen’s *d* = 3.53 for LL-SS). We also found that the variation across replicates was not significantly different across the three population types (pairwise analysis using the Fligner-Killeen test: *P* = 0.576 (LL-SL); *P* = 0.644 (LL-SS); *P* = 0.762 (SL-SS). Taken together, it is clear that EA, a major fitness-affecting trait in the presence of multiple antibiotics, had diminished in LL but enhanced in SL and SS.

**Fig. 3.**
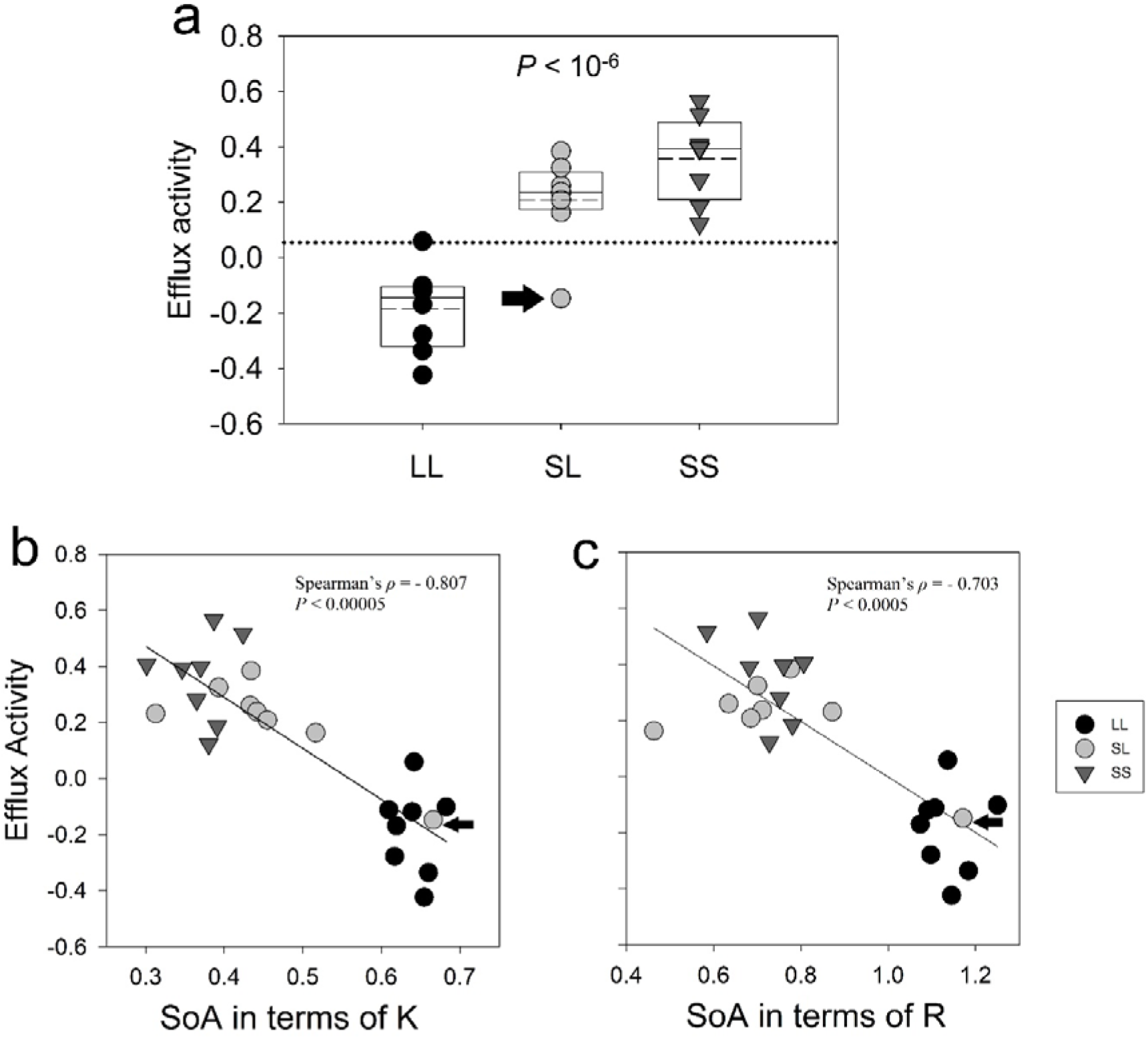
Evolved efflux activity and its correlation with the speed of adaptation. (a) EA in the three population types after evolution in the presence of the antibiotic cocktail. The solid lines in the box-plots mark the 25^th^, 50^th^, and 75^th^ percentiles; the dashed lines within the box-plots represent means (N=8). The black dotted line represents the ancestral efflux activity. Each data point represents the average of two independent efflux measurements. The grey data point marked with an arrow in each of the three panels represents the only non-LL population that lost the ancestral efflux activity (see the text for details), and is the same replicate that was an outlier in Fig 1. EA had a strong negative correlation with SoA, expressed in terms of (b) carrying capacity (K) and (c) maximum growth rate (R), respectively.

### EA was negatively correlated with SoA

A corollary of the above observations was a strong negative correlation between EA and SoA, both in terms of *K* (Fig. 3b; Spearman’s ρ = −0.807; *P* = 1.898×10^-6^) and *R* (Fig. 3c; Spearman’s ρ = −0.703; *P* = 1.256×10^-4^). Since the three population types differ in terms of their population sizes, we also checked if EA was also negatively correlated with the harmonic mean population size and found the same (Spearman’s ρ = −0.766; *P* = 1.274×10^-5^).

### LL populations fare worse in alternative environments

In the alternative environments, we found a significant population type × environment interaction, both in terms of carrying capacity (K, mixed-model ANOVA: *P* = 3.206×10^-13^; F_6,255_ = 13.42; partial η^2^ = 0.24 (large effect)) and maximum growth rate (R, mixed-model ANOVA: *P* = 3.61×10^-9^; F_6,255_ = 9.244; partial η^2^ = 0.178 (large effect)). As the interaction terms were significant, we chose not to interpret the main-effects.

To determine the effects of population type on fitness in alternative environments, we performed a separate analysis for each environment, and subjected it to a stringent set of conditions before establishing statistical significance (see Materials and methods). We found that amongst the three population types, LL had the lowest fitness in alternative environments. In terms of K, LL had the lowest fitness in all four alternative environments (Fig. 4a). In terms of R, LL had the lowest fitness in two alternative environments (Fig. 4b). Importantly, there was no environment in which LL had significantly higher fitness than SL or SS. As with EA, the observations made in Fig. 4 can only be attributed to differences in the population sizes of LL, SL, and SS during evolution in the antibiotic cocktails.

**Fig. 4.**
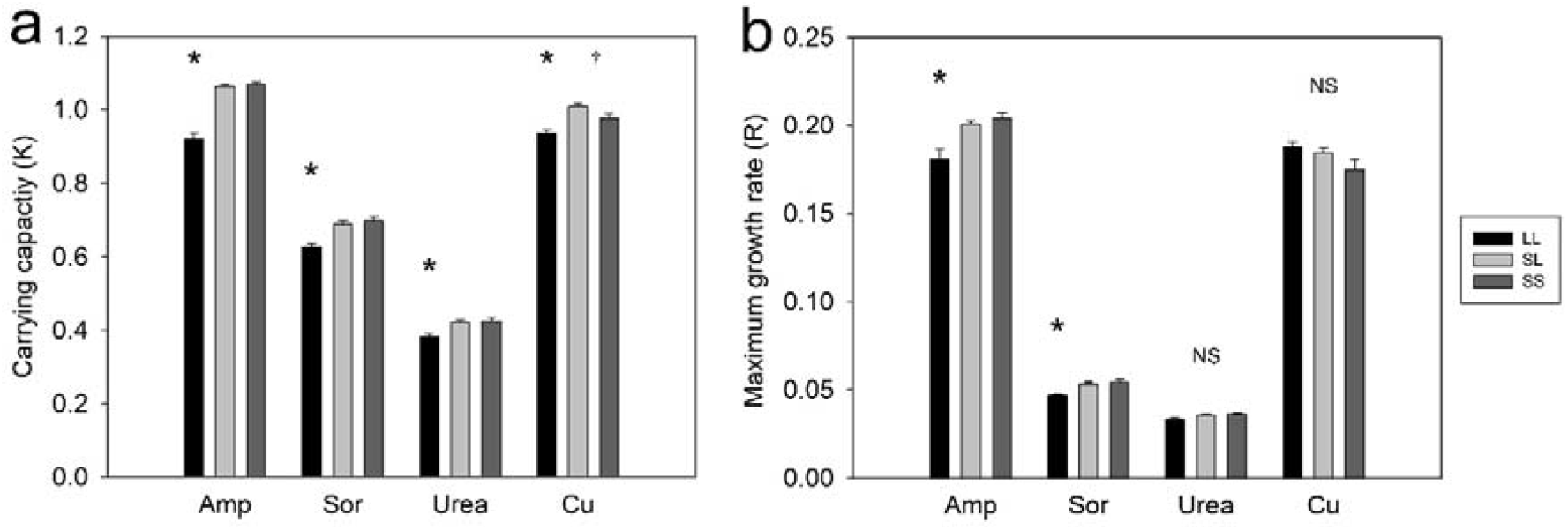
LL had significantly lower fitness than SL and SS in alternative environments. (a) Fitness expressed in terms of K (mean ± SEM; N = 8). (b) Fitness expressed in terms of R (mean ± SEM; N = 8). * refers to cases where the following four conditions are met simultaneously: (1) The ANOVA for the population type triplet reveals significant differences after the Benjamini-Hochberg procedure. (2) The difference within the population type triplet has large effect size (partial η^2^). (3) The pairwise differences between LL-SL and LL-SS are significant (Tukey’s HSD (post-hoc)). (4) The pairwise differences between LL-SL and LL-SS have large or medium effect sizes (Cohen’s d). 11 out of 12 pairwise differences marked by * had large effect sizes. † refers to the only case where SL was significantly different from SS. ‘NS’ refers to cases where the ANOVA for the population type triplet reveals no significant differences after Benjamini-Hochberg procedure. See Table S1 for detailed statistical results.

## Discussion

### Large populations pay a cost for adapting faster

The three population types (LL, SL, and SS) had experienced identical selection environments containing the antibiotic cocktail. An earlier study had reported that there was no significant difference in their fitness in the selection-environment at the same time point at which EA was measured in this study (Chavhan et al. 2019). Moreover, all the three population types had descended from the same ancestral colony. Therefore, the observations in Fig. 2 and Fig. 3 can only be attributed to the differences in their population sizes experienced during their selection history.

The extant literature shows that increased EA can improve the performance of *E. coli* in the presence of the antibiotics used in our study (Morita et al. 1998; Nishino et al. 2009; Nikaido and Pagès 2012). Hence, the presence of the antibiotics in the selection environment would intuitively suggest that EA should either be conserved or enhanced during adaptation to this environment. This is consistent with the increase in EA in the SL and SS lines (Fig. 3). However, the decay of EA in LL lines demonstrates that even under the same selection environment, whether a fitness-related trait will enhance or decay, can depend on the population size faced during selection.

The role of population size in affecting the evolution of a trait is extremely well studied since the days of Sewall Wright (Goodhart 1963; Wright 1984; Charlesworth 2009). All else being equal, for a given magnitude of stress, larger populations entail reduced effects of drift and therefore, stronger effects of selection (Charlesworth 2009). All the experimental populations in our study were large enough for their evolutionary dynamics to be driven primarily via selection and not by drift (Desai et al. 2007; Sniegowski and Gerrish 2010; Cooper 2018).

### Loss of efflux activity is primarily due to pleiotropic response

Evolutionary changes in a biological character like EA can be explained by two mechanisms that need not be mutually exclusive (Cooper and Lenski 2000; Dorken et al. 2004; Maughan et al. 2006; Hall and Colegrave 2008). The first of these is the accumulation of mutations that are neutral to fitness in the selection environment but non-neutral to the biological character in question (conventionally known as mutation accumulation (MA) (Kimura 1983; Kawecki et al. 1997; Cooper and Lenski 2000). The other mechanism is pleiotropy, in which the adaptive variation which gets selected in the selection environment affects the biological character in question non-neutrally (Rose and Charlesworth 1980; Cohan et al. 1994; Holt 1996; Cooper 2014). However, MA is unlikely to play a significant role in the evolution of characters that undergo experimentally detectable phenotypic changes within a few hundred generations (Kassen 2002; Cooper 2018) (see Supplementary Text for more discussion).

Pleiotropic responses are expected to be correlated with both the speed of adaptation (SoA) and population size. We found strong negative correlations between EA and SoA (Fig. 3 (b and c)), as well as between EA and the harmonic mean of population size. Furthermore, we found that the only non-LL population that lost its ancestral EA (SL - replicate 4, the outlier marked with an arrow in Figs. 2 and 3) was also the only outlier in terms of SoA. This outlier was similar to LL populations in terms of adaptation-speed. In other words, SL – replicate 4 was an outlier in terms of EA and SoA. However, it was not an outlier in terms of the negative correlations shown in Fig 3, making the correlations stronger. Thus, our experimental design enabled us to attribute the differences in population size across our treatments to the differences in the pleiotropic responses which shaped the evolution of EA (antagonistic pleiotropy in LL but synergistic pleiotropy in SL and SS).

### Quantitative differences in population size can lead to qualitative differences in the result of selection on characters

Biological characters can be lost over evolutionary time if they are unessential or disadvantageous (Fong et al. 1995; Porter and Crandall 2003; Jeffery 2005; Visser et al. 2010). The fate of the character in question (whether it decays, gets maintained, or enhances) during evolution is expected to be determined by the environment in which evolution occurs (Cooper 2014). For example, extremely dark environments have been invoked to explain the loss of eyes in multiple systems (Jeffery 2005; Protas et al. 2011). Similarly, the metabolic erosion observed over >50,000 generations in the Lenski Long-Term Evolution Experiment (LTEE) has been linked to the presence of only one usable carbon source throughout evolution (Leiby and Marx 2014). Moreover, the environment in which evolution happens is conventionally assumed to be the major factor in determining the utility of the biological character in question (Hall and Colegrave 2008). If in a given environment, the evolution of the biological character is largely affected by selection and not by drift, it is expected to result either in enhancement/maintenance or in decay of the character (but not both). It has been demonstrated empirically that if the selection-environments are different, the same biological character can decay by disparate evolutionary mechanisms (MA versus pleiotropy) (Hall and Colegrave 2008). Our study adds to the understanding of evolutionary decay of characters by showing how the character in question can decay or enhance in the same environment based on population size (Fig 3). In other words, our study shows that the selection-environment cannot always explain divergent evolutionary fates of a biological character. An important goal of experimental evolution is to understand how quantitative differences in population genetic parameters can lead to qualitatively different evolutionary outcomes. Our study takes a step in this direction by demonstrating how quantitative differences (in population size) can translate into qualitative differences (decay or enhancement) in a fitness-related trait during evolution.

One important question to ask here is why did the efflux activity decline in the LL populations? Although it is not possible to answer this question definitively from our data, we provide a brief speculation in this regard. Efflux is known to be an energetically expensive process (Nikaido 1994). In the presence of mutations that directly reduce the effects of antibiotic(s), efflux enhancing mutations are expected to be deleterious. However, in the absence of such mutations, efflux enhancing mutations are expected to be beneficial. Hence, once mutations that directly make the antibiotic ineffective arise, decay of efflux could be beneficial to fitness. Owing to their large population size, the LL populations could have accessed rare large-effect beneficial mutations for loci that directly render the antibiotic(s) ineffective. Therefore, once such mutations arose, the LL populations could increase their fitness by the decay of efflux. On the other hand, Inaccessibility to such rare mutations in the small populations (SL and SS) could have led to positive selection for efflux, which manifested as enhanced EA levels in SL and SS after ∼380 generations. We note that recent advances in whole-genome whole population sequence analysis coupled with appropriate genetic manipulation can potentially be used to validate these speculations (Long et al. 2015; Anand et al. 2016; Cooper 2018). However, such an analysis is outside the scope of the present study.

### Large populations had lower fitness in alternative environments

The LL populations not only evolved reduced EA (Fig. 3), they also had the lowest fitness among the three types in alternative environments (Fig. 4). It should be noted that the reduced EA of LL can be linked to low fitness only in two of the four alternative environments (ampicillin and high [Cu^2+^]). This is because the other two alternative environments challenged the bacterial population with nutrient-poor conditions (not with xenobiotic chemicals), where EA is not expected to directly provide a fitness advantage.

In this study, we use random samples of potentially heterogeneous populations for assaying fitness in the alternative stressful environments. Such assays can potentially be driven by rare outliers with very high fitness values under these unexplored conditions, leading to inflated estimates of mean population fitness. We find that in the alternative stressful environments, the larger populations (LL) had lower fitness than the smaller ones (SL and SS). If the observed population-level fitness of LL was driven largely by the rare high fitness genotypes, when we remove the effects of these outliers, the robust (i.e., outlier-removed) value of the mean phenotype of the LL populations would be even lower than what is currently reported. This implies that our estimates about the reduction in fitness of LLs (vis-à-vis the SL and SS populations) is conservative. Technically, the same argument could work for the SL and SS populations too. However, since the size of these populations is ∼16,500 times less than the LL populations, theoretically one expects the SL and SS populations to have relatively milder outliers owing to their low supply of variation. Thus, our estimates of fitness are expected to be more inflated in the larger populations as compared to the smaller ones. Importantly, it is highly unlikely that the inflation is higher for the smaller populations. Thus, our observations regarding fitness in the alternative environments are likely to be robust to the potential effects of rare highly fit outliers.

The fitness trajectories of asexual populations are known to show a decrease in the rate of fitness-increase as adaptation progresses (Cooper and Lenski 2000; Elena and Lenski 2003; Couce and Tenaillon 2015; Tenaillon et al. 2016; Chavhan et al. 2019). If pleiotropy plays a major role in shaping fitness in alternative environments, periods of faster adaptation are known to cause greater loss of unused functions within in a given population (Cooper and Lenski 2000). This leads to the expectation that larger populations (which adapt faster) should have lower fitness in alternative environments. We found that this indeed was the case as the LL populations had lower fitness in the alternative environments in general. Our results are applicable across populations of different sizes and are congruent with those of Cooper and Lenski (2000) whose study was applicable within single populations at different times during their evolution. However, since the three population types in our experiments eventually reached similar fitness in the selection-environment despite initially adapting at different speeds, theory predicts that simple pleiotropic responses arising from beneficial mutations should lead to similar fitness across the three types in the alternative environments. Since the three population types had significantly different fitness in the alternative environments, such simple pleiotropy cannot possibly explain our results.

The theory of asexual adaptive dynamics predicts that in a population of effective size ‘N_e_’ and rate of spontaneous beneficial mutations (per genome per cell division) ‘U_b_’, the number of beneficial mutations that can potentially compete with a beneficial mutation of selection coefficient ‘s’ on its way to fixation is given by 2N_e_U_b_ln(N_e_s/2) (Sniegowski and Gerrish 2010). For all the three population types, even with highly conservative estimates of U_b_ = 0.0001 and s = 0.01, this number amounts to more than a thousand competing beneficial mutations (the harmonic mean sizes of our experimental populations were close to 10^10^ for LL, and 10^6^ for SL and SS). Moreover, both the measures of fitness (growth rate and carrying capacity) had increased by more than 1.5-fold in all the experimental populations by the end of the experiment (Chavhan et al. 2019). This suggests that the reference value of ‘s’ should be much larger than 0.01, making the number of competing beneficial mutations even larger. A previous ∼500 generations long experimental evolution study had found that asexual yeast populations adapted via dynamics that were best explained by the multiple-mutations paradigm (Desai et al. 2007). This paradigm implies that multiple beneficial mutations occurring within a lineage can simultaneously rise to high frequencies (Desai and Fisher 2007; Desai et al. 2007). The effective size (i.e., harmonic mean ∼10^6^) of even the smallest populations in our study (SL/SS) were similar to the largest population in Desai et al. (2007). Thus, for the given evolutionary time scale and population sizes, the adaptive events in our experimental populations are likely to be based on multiple mutations. However, the observations regarding fitness in the alternative environments cannot still be explained using a simple (i.e. linear / additive) combination of pleiotropic effects of multiple beneficial mutations. This is because if pleiotropic effects of multiple mutations just combined additively then, given that all three population types (i.e., LL/SL/SS) had the same fitness in the selection environments, they would have shown similar fitness values in the alternative environments too. However, this was not the case (Fig 4). Thus, to explain the fitness patterns in alternative environments, one requires further assumptions about how the pleiotropic effects of multiple mutations interact with each other.

One such assumption can be that the pleiotropic effects of beneficial mutations increase more rapidly than linear (say exponential or any other similar non-linear function) with the magnitude of direct effects. This is similar to the key assumption made by previous studies that large beneficial mutations have substantial pleiotropic effects while small beneficial mutations show negligible pleiotropy (Lande 1983). Moreover, the pleiotropic effect of a combination of multiple mutations can be smaller than the sum of their individual pleiotropic effects, as found by a previous study on *Escherichia coli* populations (Bohannan et al. 1999). Finally, we note that these assumptions/possibilities are not mutually exclusive, and their simultaneous action can also explain our observations.

Unfortunately, few studies have rigorously investigated the pleiotropic effects stemming from a combination of multiple mutations (Flynn et al. 2013; Schick et al. 2015). A recent study demonstrated that although the direct fitness effects of combinations of mutations consistently showed diminishing returns in the selection-environments, the pleiotropic fitness effects of such mutational combinations were highly variable in alternative environments (Schick et al. 2015). In other words, the pleiotropic fitness effects of combination of mutations were less than the sum of individual pleiotropic effects in some cases but more than the sum in others. (Schick et al. 2015). Unfortunately, we are not in a position to ascertain the exact nature of the relationship between pleiotropy and fitness, a predicament succinctly summed up by Cooper (2014): “The uncertainty of the form of pleiotropic effects reflects a general lack of understanding of how mutations interact to affect fitness, particularly over the long term.”

In summary, our observations regarding the performance in alternative environments suggest that pleiotropy can potentially explain the link between SoA and preparedness for alternative environmental conditions, and mechanisms like EA need not always be invoked.

The idea that large populations can adapt so effectively and rapidly to their constant environment that they can be rendered vulnerable to future environmental change is new and counterintuitive. Our results could be relevant for understanding evolution in asexual populations experiencing changes between environments with fitness trade-offs (de Roode et al. 2008; Bahri et al. 2009; Andersson and Hughes 2010). Our study shows that large populations can have lower fitness in alternative environments (Fig. 4). In such populations, adaptation is expected to be driven by mutations with large fitness benefits (Sniegowski and Gerrish 2010), which are typically assumed to be associated with heavier pleiotropic disadvantages (Lande 1983; Orr and Coyne 1992) that can lead to greater fitness trade-offs. Thus, a logical next step would be to test a putative relationship between population sizes and fitness-tradeoffs using reciprocal selection experiments in multiple environments. This can lead to a better understanding of the population genetics of fitness trade-offs and ecological specialization (Fry 1996; Cooper and Lenski 2000; Kassen 2002, 2014; Rodríguez-Verdugo et al. 2014; Schick et al. 2015).

It is a well-established notion that very small population sizes can lead to such strong effects of genetic drift that the latter can overshadow selection and preclude adaptation (Charlesworth 2009). Here we show that very large population sizes (as in LL), while leading to rapid adaptation in the current environment, can also render populations vulnerable to sudden environmental changes. Taken together, these insights point to a trade-off between maximizing adaptation rate and avoiding becoming vulnerable to environmental changes. Thus, populations that are small enough to avoid pleiotropic disadvantages but large enough to adapt (albeit slowly) to the current conditions (like SL and SS) can face environmental changes better than very large populations (like LL). We have used periodically bottlenecked bacterial populations to demonstrate the above trade-off. Therefore, our counterintuitive results can potentially have important implications for the fate of naturally occurring microbial populations that face periodic bottlenecks (e.g., host-to-host transfer of gut microbiota or pathogens), particularly if their environment changes in bouts. By demonstrating a novel link between population size and the immediate response to sudden environmental changes, our study thus adds to the prospering field of the evolution of evolvability (Carter et al. 2005; Jones et al. 2007; Crombach and Hogeweg 2008; Wagner 2013).

## Supplementary information

### A. Supplementary Methods

#### 1. Composition of the selection medium

Composition of Nutrient Broth (Himedia Laboratories Pvt. Ltd.):

- Peptic digest of animal tissue (5 g/l)
- Sodium chloride (5 g/l)
- Beef extract (1.50 g/l)
- Yeast extract (1.50 g/l)

We added the following cocktail of antibiotics to the Nutrient Broth:

1. Norfloxacin (0.015 μg/ml)
2. Rifampicin (6 μg/ml)
3. Streptomycin (0.1 μg/ml)

#### 2. Details of the protocol used for measuring energy-dependent efflux activity

We used a previously-established fluorescence-based assay for measuring energy-dependent efflux activity (EA) in Gram negative bacteria (Webber and Coldham 2010; Karve et al. 2015). We used a small molecule (bis-benzimide) that enters bacterial cells and fluoresces after intercalating with DNA (bis-benzimide excites at 355 nm and emits at 465 nm). The details of the protocol are as follows:

- Cryostocks belonging to each of the 25 populations (24 descendants and 1 ancestor) were revived in Nutrient Broth (NB) for 18 hours.
- The revived populations were brought to similar sizes by dilutions with (NB) so that the OD at 600 nm was between 0.03 and 0.06 when measured using Nanodrop (Thermo Scientific, 2000).
- The above cultures (with similar sizes) were centrifuged at 13,400 rpm for 2 minutes. The supernatant was discarded, and PBS buffer (at pH 7.4) was used to resuspend the pellet.
- A small aliquot of the ancestral culture was heated at 60 □ C to set the upper range of fluorescence gain.
- 8 μl of glucose solution (1% w/v) was also added because the aim was to measure energy-dependent efflux activity.
- 20 μl of bis-benzimide was added to each well containing 168 μl live culture. The same volume of bis-benzimide was also added to the control well with dead cells. An automated plate reader (Tecan Infinite M200 Pro) was used to measure fluorescence. Initially, we measured fluorescence for 40 min. All the fluorescence curves had reached a steady state by 35 min.
- After 40 minutes, the plate was taken out and 4 μl of a non-specific inhibitor of active efflux was added to all the wells. The inhibitor of efflux used in our study was CCCP (Carbonyl Cyanide m-Chlorophenylhydrazone; C2759 Sigma).
- The measurement of fluorescence was resumed and continued for a further 30 minutes so that a new steady state could be reached by all the fluorescent curves.
- Efflux activity was measured using the following formula: (Fluorescence at 70 min – Fluorescence at 35 min) / (Fluorescence at 35 min)

## Supplementary Results

### 1. Statistical details of ANOVAs done separately in each alternative environment

**Table S1.**
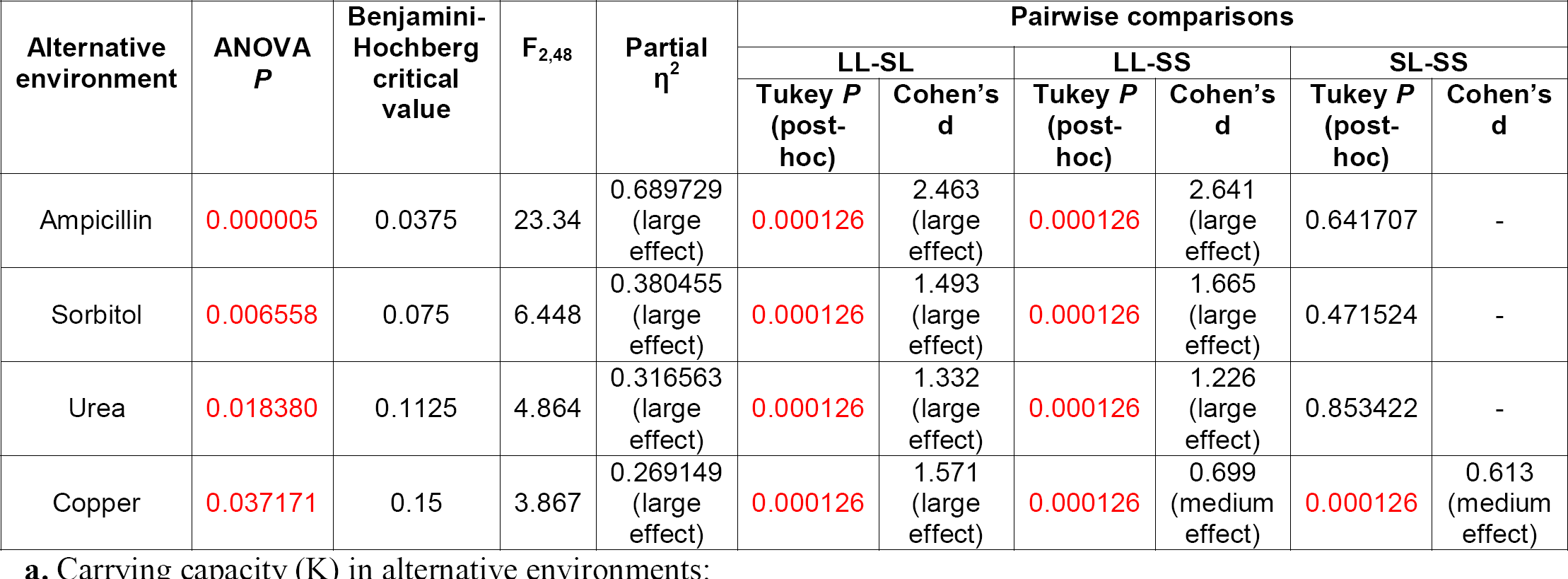
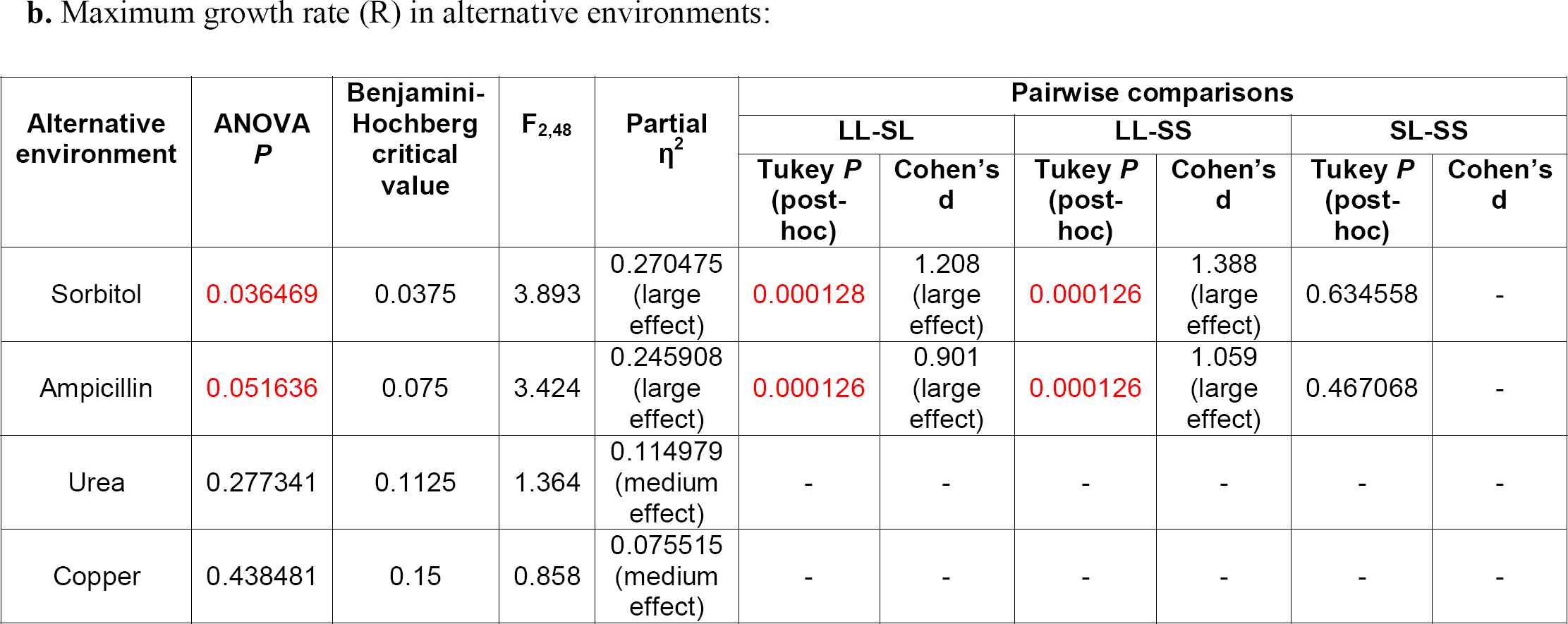
Summary of the statistical analysis done separately for each of the four alternative environments. **(a)** Analysis in terms of K. **(b)** Analysis in terms of R. Statistically significant P values are shown in red. The False Discover Rate (FDR) used here is 0.15 (McDonald 2009). Partial η^2^ was interpreted as: Partial η^2^ < 0.06 (small effect); 0.06 < Partial η^2^ < 0.14 (medium effect); 0.14 < Partial η^2^ (large effect). Cohen’s d was interpreted as: 0.2 < d < 0.5 (small effect), 0.5 < d < 0.8 (medium effect); d > 0.8 (large effect). Tukey’s post-hoc test was done only when the ANOVA results were significant after the Benjamini Hochberg procedure. Cohen’s d was interpreted only when the pairwise differences (revealed by Tukey’s post-hoc test) were significant.

1. Statistical details of ANOVAs done separately in each alternative environment:
| Alternative environment | ANOVA *P* | Benjamini-Hochberg critical value | $F_{2,48}$ | Partial $\eta^2$ | Pairwise comparisons | | | | | |
| --- | --- | --- | --- | --- | --- | --- | --- | --- | --- | --- |
|  |  |  |  |  | LL-SL |  | LL-SS |  | SL-SS |  |
|  |  |  |  |  | Tukey *P* (post-hoc) | Cohen's *d* | Tukey *P* (post-hoc) | Cohen's *d* | Tukey *P* (post-hoc) | Cohen's *d* |
| Ampicillin | 0.000005 | 0.0375 | 23.34 | 0.689729 (large effect) | 0.000126 | 2.463 (large effect) | 0.000126 | 2.641 (large effect) | 0.641707 | - |
| Sorbitol | 0.006558 | 0.075 | 6.448 | 0.380455 (large effect) | 0.000126 | 1.493 (large effect) | 0.000126 | 1.665 (large effect) | 0.471524 | - |
| Urea | 0.018380 | 0.1125 | 4.864 | 0.316563 (large effect) | 0.000126 | 1.332 (large effect) | 0.000126 | 1.226 (large effect) | 0.853422 | - |
| Copper | 0.037171 | 0.15 | 3.867 | 0.269149 (large effect) | 0.000126 | 1.571 (large effect) | 0.000126 | 0.699 (medium effect) | 0.000126 | 0.613 (medium effect) |

**Table S1. Summary of the statistical analysis done separately for each of the four alternative environments. (a) Analysis in terms of K. (b) Analysis in terms of R.**
| Alternative environment | ANOVA *P* | Benjamini-Hochberg critical value | $F_{2,48}$ | Partial $\eta^2$ | Pairwise comparisons | | | | | |
| --- | --- | --- | --- | --- | --- | --- | --- | --- | --- | --- |
|  |  |  |  |  | LL-SL |  | LL-SS |  | SL-SS |  |
|  |  |  |  |  | Tukey *P* (post-hoc) | Cohen's *d* | Tukey *P* (post-hoc) | Cohen's *d* | Tukey *P* (post-hoc) | Cohen's *d* |
| Sorbitol | 0.036469 | 0.0375 | 3.893 | 0.270475 (large effect) | 0.000128 | 1.208 (large effect) | 0.000126 | 1.388 (large effect) | 0.634558 | - |
| Ampicillin | 0.051636 | 0.075 | 3.424 | 0.245908 (large effect) | 0.000126 | 0.901 (large effect) | 0.000126 | 1.059 (large effect) | 0.467068 | - |
| Urea | 0.277341 | 0.1125 | 1.364 | 0.114979 (medium effect) | - | - | - | - | - | - |
| Copper | 0.438481 | 0.15 | 0.858 | 0.075515 (medium effect) | - | - | - | - | - | - |

### 2. Ancestral fitness values in the alternative environments

**Table S2.**

| Alternative environment | K | R |
| --- | --- | --- |
| Sorbitol | 0.652 | 0.054 |
| Ampicillin | 0.836 | 0.118 |
| Urea | 0.427 | 0.049 |
| Copper | 0.924 | 0.170 |

## Supplementary Text

### Why mutation accumulation via drift cannot explain the evolution of efflux activity

As discussed in the main-text, all three population types were derived from the same ancestor. Importantly, all the three types had such large population sizes that drift over a few hundred generations is not expected to produce phenotypically detectable changes (Desai and Fisher 2007; Sniegowski and Gerrish 2010; Cooper 2018). In other words, a time period of approximately 380 generations is too short to observe significant effects of MA in our experiments. However, some previous studies have attributed phenotypic changes in trait values to MA over similar time-scales in similarly sized populations (Collins and Bell 2004; Hall and Colegrave 2008). Therefore, we briefly examined whether MA can have a significant effect on the evolution of efflux activity in our populations.

Mutation accumulation (MA) and pleiotropy have contrasting dependencies on a variety of population genetic parameters. MA is positively related with the rate of spontaneous mutations per individual per generation (µ) (Kimura 1983; Hall and Colegrave 2008), but is independent of the population size (N) (Kimura 1983) and the speed of adaptation (Cooper and Lenski 2000; Hall and Colegrave 2008; Cooper 2014). On the other hand, pleiotropic responses are influenced by both N and µ (Kimura 1983; Hall and Colegrave 2008), and are expected to be correlated with the speed of adaptation (SoA) (Cooper and Lenski 2000; Cooper 2014).

In the main-text, we establish why the evolution of efflux activity can be largely explained using pleiotropy. In contrast to the strong correlations reported in Fig. 3 (b and c) of the main text, EA is not expected to be correlated with either SoA or N if efflux evolves via MA.

If the LL populations had attained much higher mutation rates than SL and SS during ∼ 380 generations of evolution in identical environments, then they could be expected to show the lowest EA due to heaviest MA. However, any mutation that could increase µ in LL would have to appear *de novo* with the very small frequency of 1/N. Since selection does not act directly on mutation-rate-altering loci (Chao and Cox 1983; Sniegowski et al. 1997; Orr 2000; Gentile et al. 2011), µ-altering mutations spread via hitchhiking with mutations that are direct targets of selection. Thus, a mutation that increases µ (henceforth referred to as ‘mutator’) can only rise via random drift before an extra non-neutral mutation happens in its lineage. Since the occurrence of two mutations within a lineage is a highly improbable event (the maximum probability being µ^2^), the mutator needs to rise to large frequencies before it can start hitchhiking with a non-neutral mutation. This makes the establishment of the mutator highly unlikely if N is large. It has been demonstrated that mutators rise successfully via hitchhiking only if their initial frequency is larger than a threshold (Chao and Cox 1983). Indeed, *de novo* mutators have been shown to go to extinction in most replicate populations (Sniegowski et al. 1997, 2000; Taddei et al. 1997; Giraud et al. 2001; De Visser and Rozen 2005; Raynes and Sniegowski 2014). Therefore, MA is unlikely to explain the fact that almost all replicates of SL and SS increased EA while all LL replicates had evolved reduced EA.

Furthermore, among populations that are large enough to experience clonal interference, the evolution of high mutation rate, leading to greater MA, is more likely to happen in smaller populations and not in larger ones (De Visser and Rozen 2005; Desai and Fisher 2007; Raynes et al. 2012). Indeed, recent experimental studies have demonstrated that despite starting with initial frequencies as high as 30%, the variants with higher mutation rates go to extinction in large asexual populations but hitchhike to fixation in small ones (Gentile et al. 2011; Raynes et al. 2012). Thus, even in the unlikely scenario of MA influencing efflux over a time-scale of a few hundred generations, efflux was more likely to decay in SL and SS, not in LL. Since we observed the decay of efflux in LL and not SL or SS (Fig. 3), MA is unlikely to be an explanation for the observed EA patterns.

## Acknowledgements

We thank Milind Watve and MS Madhusudhan for their valuable inputs. YDC was supported by a Senior Research Fellowship initially sponsored by IISER Pune and then by Council for Scientific and Industrial Research (CSIR), Govt of India. This project was supported by an external grant (BT/PR5655/BRB/10/1088/2012) from Department of Biotechnology, Govt of India, and internal funding from IISER Pune.

